# Multiple adjoining word- and face-selective regions in ventral temporal cortex exhibit distinct dynamics

**DOI:** 10.1101/2020.12.29.424760

**Authors:** Matthew J. Boring, Edward H. Silson, Michael J. Ward, R. Mark Richardson, Julie A. Fiez, Chris I. Baker, Avniel Singh Ghuman

**Affiliations:** Center for Neuroscience at the University of Pittsburgh, University of Pittsburgh, A210 Langley Hall, Pittsburgh, PA 15260, USA; Center for the Neural Basis of Cognition, 4400 Fifth Avenue, Pittsburgh, PA 15213; Department of Neurological Surgery, University of Pittsburgh Medical Center, 200 Lothrop Street, Pittsburgh, PA 15213, USA; National Institute of Mental Health, National Institutes of Health, Magnuson Clinical Center, Bethesda, MD 20814 USA; School of Philosophy, Psychology and Language Sciences, University of Edinburgh, 7 George Square, Edinburgh EH8 9JZ, UK; Department of Neurosurgery, Massachusetts General Hospital, 55 Fruit Street, Boston, MA 02144, USA; Harvard Medical School, 25 Shattuck St., Boston, MA 02115, USA; Department of Psychology, University of Pittsburgh, 210 South Bouquet St, Pittsburgh, PA 15260, USA

## Abstract

The map of category-selectivity in human ventral temporal cortex (VTC) provides organizational constraints to models of object recognition. One important principle is lateral-medial response biases to stimuli that are typically viewed in the center or periphery of the visual field. However, little is known about the relative temporal dynamics and location of regions that respond preferentially to stimulus classes that are centrally viewed, like the face- and word-processing networks. Here, word- and face-selective regions within VTC were mapped using intracranial recordings from 36 patients. Partially overlapping, but also anatomically dissociable patches of face- and word-selectivity were found in VTC. In addition to canonical word-selective regions along the left posterior occipitotemporal sulcus, selectivity was also located medial and anterior to face-selective regions on the fusiform gyrus at the group level and within individual subjects. These regions were replicated using 7 Tesla fMRI in healthy subjects. Left hemisphere word-selective regions preceded right hemisphere responses by 125 ms, potentially reflecting the left hemisphere bias for language; with no hemispheric difference in face-selective response latency. Word-selective regions along the posterior fusiform responded first, then spread medially and laterally, then anteriorally. Face-selective responses were first seen in posterior fusiform regions bilaterally, then proceeded anteriorally from there. For both words and faces, the relative delay between regions was longer than would be predicted by purely feedforward models of visual processing. The distinct time-courses of responses across these regions, and between hemispheres, suggest a complex and dynamic functional circuit supports face and word perception.

**Significance Statement:** Representations of visual objects in the human brain have been shown to be organized by several principles, including whether those objects tend to be viewed centrally or peripherally in the visual field. However, it remains unclear how regions that process objects that are viewed centrally, like words and faces, are organized relative to one another. Here, invasive and non-invasive neuroimaging suggests there is a mosaic of regions in ventral temporal cortex that respond selectively to either words or faces. These regions display differences in the strength and timing of their responses, both within and between brain hemispheres, suggesting they play different roles in perception. These results illuminate extended, bilateral, and dynamic brain pathways that support face perception and reading.

## Introduction

Investigations into the spatial organization of category-selectivity in ventral temporal cortex (VTC) have been instrumental in establishing several organizational principles of the visual system. Functional magnetic resonance imaging (fMRI) studies have helped identify lateral-medial biases in ventral stream responses to objects depending on where they typically appear in the visual field (retinotopic eccentricity) (Hasson et al., 2002; Konkle and Caramazza, 2013; Grill-Spector and Weiner, 2014). Specifically, lateral regions of VTC are selective for objects that tend to be viewed centrally (foveated), like words and faces, whereas more medial regions are selective for objects that tend to fall on the periphery of the retina, like navigationally relevant information such as buildings (Haxby et al., 1996; Aguirre et al., 1998; Cohen et al., 2000; Hasson et al., 2002). This broad principle of organization by eccentricity fails to inform us about how representations of different stimuli that are foveated, like words and faces, are organized in VTC relative to one another.

Despite sharing similar typical retinotopic eccentricity, word and face stimuli are highly distinct along several axes that are also hypothesized to influence where they are processed in VTC (Op de Beeck et al., 2019). Word- and face-processing operate on very different low-level visual properties (Kay and Yeatman, 2017), follow different developmental trajectories (Saygin et al., 2016), and feed into distinct networks that support either language or social interactions (Stevens et al., 2015, 2017), respectively. Despite this, the cortical localizations for word- and face-processing in VTC are remarkably close together, and it remains debated whether or not there are regions in VTC that independently encode word or face information at all (Behrmann and Plaut, 2013).

Neuroimaging studies have separately mapped word- and face-processing networks in VTC. Printed word recognition is thought to be carried out in part by a network of regions along the left occipitotemporal sulcus, that differ in the complexity of their responses and are thought to be hierarchically organized (Halgren et al., 1994; Cohen et al., 2000; Vinckier et al., 2007; Dehaene and Cohen, 2011; Lerma-Usabiaga et al., 2018). Face-processing is thought to be carried out in part by a network of regions distributed bilaterally along the midfusiform sulcus (Tsao et al., 2008; Weiner and Grill-Spector, 2010). However, few studies have investigated VTC’s responses to word and face stimuli within the same participants (Allison et al., 1994; Haxby et al., 1994; Puce et al., 1996; Matsuo et al., 2015; Harris et al., 2016). Those that have, have relied on low sample sizes or imaging modalities with differential sensitivity to different aspects of neural activity (like high and low-frequency neural activity (Engell et al., 2012; Jonas et al., 2016)). Therefore, much remains unknown about how visual word- and face-processing networks organize relative to one another, and to what degree they overlap (Haxby et al., 1994; Puce et al., 1996; Dehaene et al., 2010; Matsuo et al., 2015; Harris et al., 2016).

Further, word- and face-selective regions have primarily been mapped using methods lacking high spatiotemporal resolution. Therefore, it is unclear if the nodes within these processing networks differ in the temporal dynamics of their responses, although previous studies have suggested that different regions may contribute to distinct stages of word- and face-processing (Federmeier and Kutas, 1999; Vinckier et al., 2007; Li et al., 2018). Further, category-selective maps derived from BOLD responses may be incomplete due to BOLD’s increased sensitivity to early stimulus evoked activity (100-300 ms after stimulus presentations) relative to later responses (Jacques et al., 2016; Ghuman and Martin, 2019) and greater correlation with high frequency broadband activity in invasive neural recordings compared to lower-frequency electrical potentials (Engell et al., 2012; Jacques et al., 2016).

In the present study, we characterized the spatial organization and functional dynamics of word- and face-processing networks within VTC using intracranial electroencephalography (iEEG) data collected from 36 patients with pharmacologically intractable epilepsy and 7 T fMRI data collected from eight healthy participants.

## Materials and Methods

### Intracranial EEG data collection and preprocessing

#### Participants

38 patients (14 males, ages 19-65, 32 righthanded) had intracranial surface and/or depth electrodes implanted for the treatment of pharmacologically intractable epilepsy. Depth electrodes were produced by Ad-Tech Medical and PMT Corporation and were 0.86 and 0.8 mm in diameter, respectively. Grid electrodes were produced by PMT Corporation and were 4 mm in diameter. Because depth electrode contacts are cylindrical, the surface area of the recording site was similar across grid and strip electrode contacts. To be concise, “electrode contacts” are referenced to as “electrodes” throughout the manuscript. No consistent differences in neural responses were observed between grid and depth electrodes. Only electrodes implanted in ventral temporal cortex, defined as below the inferior temporal gyrus and anterior to the posterior tip of the fusiform in the participant-centered space, were considered in this study. Two patients did not have any electrodes within this region of interest, therefore only data from 36 participants were analyzed for this study. Electrodes identified as belonging to the seizure onset zone based on the clinical report or showing epileptiform activity during the tasks were excluded from the analysis. All participants gave written informed consent. The study was approved by the University of Pittsburgh Institutional Review Board. Patients were monetarily compensated for their time.

Electrodes were localized via either post-operative magnetic resonance imaging (MRI) or computed tomography scans co-registered to the pre-operative MRI using Brainstorm (Tadel et al., 2011). Surface electrodes were projected to the nearest point on the pre-operative cortical surface automatically parcellated via Freesurfer (Dale et al., 1999) to correct for brainshift (Hermes et al., 2010). Electrode coordinates were then coregistered via surface-based transformations to the fsaverage template using Freesurfer cortical reconstructions.

#### Experimental Design

All participants underwent a category localizer task where they viewed grayscale images presented on a computer screen positioned two meters from their face. Images occupied approximately 6 x 6 degrees of visual angle and were presented for 900 ms with 1500 ms inter-stimulus interval with random 400 ms jitter. Participants were instructed to press a button every time an image was presented twice in a row (1/6 of the trials). These repeat trials were excluded from the analysis yielding 70 trials per stimulus category left for analysis. Several participants underwent multiple runs of this task and therefore had 140-210 trials per stimulus category.

31 of the participants saw pictures of faces, words, bodies, hammers, houses and phase-scrambled faces. The remaining participants viewed a modified set of stimuli with the same viewing parameters described above. One participant viewed pictures of consonant-strings and pseudowords instead of hammers, two viewed shoes instead of words, one viewed consonant-strings and pseudowords instead of hammers and houses, and one viewed general tools and animals instead of hammers.

A subset of the participants that underwent the category localizer task also participated in word and/or face individuation tasks (Table 1). These tasks shared identical presentation parameters as the category-localizer task (i.e. inter-stimulus interval, stimulus-on time, and viewing angle) but contained different images. Twelve underwent a word individuation task that included pictures of real words, pseudowords, and consonant-strings or false-fonts. Participants again were instructed to respond if a given stimulus was repeated twice in a row. Every stimulus (i.e. individual word) was presented sixty times. Twenty underwent a face individuation task where they viewed individuals of varying identity and emotions. Participants were instructed to indicate if each face was male or female during this task. Each identity was repeated 60 times.

**Table 1.** iEEG participant coverage

| Number | Tasks completed | Electrodes in VTC | Face-selective | Word-selective | House-selective | Word medial to face-selective | Alternating word- and face-selective |
| --- | --- | --- | --- | --- | --- | --- | --- |
| 1 | CL | L: 6 | 0 | 0 | 0 | N/A | N/A |
| 2 | CL | L: 11 | 0 | 0 | 0 | N/A | N/A |
| 3 | CL | L: 34, R: 18 | 0 | 0 | L: 2, R: 2 | N/A | N/A |
| 4 | CL | L: 20, R: 14 | 0 | 0 | R: 2 | N/A | N/A |
| 5 | CL | R: 18 | R: 2 | 0 | 0 | N/A | N/A |
| 6 (Fig. 4B) | CL | L: 11 | L: 1 | L: 1 | 0 | Yes | N/A |
| 7 | CL, WID | L: 17 | L: 2 | L: 1 | 0 | No | No |
| 8 | CL | R: 9 | 0 | 0 | R: 2 | N/A | N/A |
| 9 | CL, WID | R: 21 | 0 | R: 1 | 0 | N/A | N/A |
| 10 (Fig. 4B) | CL, WID, FID | L: 25, R: 16 | L: 2, R: 1 | L: 2 | 0 | Yes | Yes |
| 11 | CL, FID | L: 4, R: 23 | R: 5 | L: 1, R: 1 | R: 5 | Yes | Yes |
| 12 | CL, FID | R: 42 | R: 8 | R: 4 | R: 6 | Yes | Yes |
| 13 | CL, FID | L: 38 | 0 | L: 2 | L: 2 | N/A | N/A |
| 14 | CL, FID | L: 23, R: 24 | L: 2, R: 1 | 0 | L: 2, R: 2 | N/A | N/A |
| 15 | CL, FID | L: 30 | L: 1 | 0 | L: 2 | N/A | N/A |
| 16 | CL, FID | L: 23, R: 11 | 0 | L: 1 | R: 3 | N/A | N/A |
| 17 (Fig. 4B) | CL, WID, FID | L: 48 | L: 6 | L: 4 | L: 2 | Yes | Yes |
| 18 | CL, FID | L: 23 | 0 | 0 | L: 7 | N/A | N/A |
| 19 | CL | L: 4 | 0 | L: 2 | L: 2 | N/A | N/A |
| 20 | CL | L: 23 | 0 | 0 | 0 | N/A | N/A |
| 21 | CL | R: 11 | 0 | 0 | R: 1 | N/A | N/A |
| 22 | CL, WID, FID | R: 41 | 0 | R: 6 | 0 | N/A | N/A |
| 23 | CL | L: 10 | L: 1 | L: 2 | 0 | No | No |
| 24 | CL, FID | L: 26, R: 25 | L: 3, R: 1 | R: 2 | R: 1 | Yes | Yes |
| 25 | CL, WID, FID | L: 21, R: 19 | 0 | L: 6, R: 1 | 0 | N/A | N/A |
| 26 | CL | L: 21, R: 28 | L: 2 | L: 3 | R: 3 | No | No |
| 27 | CL, FID | L: 5, R: 18 | 0 | L: 1, R: 5 | R: 3 | N/A | N/A |
| 28 (Fig. 4A) | CL, WID, FID | L: 55 | L: 6 | L: 4 | 0 | Yes | Yes |
| 29 | CL, FID | L: 42 | L: 2 | L: 2 | 0 | Yes | No |
| 30 | CL, FID | L: 26, R: 28 | L: 1, R: 2 | R: 1 | L: 2, R: 1 | Yes | No |
| 31 | CL, WID, FID | L: 19, R: 36 | L: 1, R: 6 | 0 | R: 2 | N/A | N/A |
| 32 | CL, WID | L: 10, R: 34 | L: 1 | 0 | L: 1, R: 3 | N/A | N/A |
| 33 | CL, WID, FID | L 39, R 50 | 0 | L: 4 | L: 3, R:2 | N/A | N/A |
| 34 | CL, WID | L 24, R: 27 | R: 2 | L: 5, R: 2 | L: 1, R: 3 | No | No |
| 35 | CL, FID | L: 116 | L: 16 | L: 8 | L: 6 | Yes | Yes |
| 36 | CL, WID, FID | L: 129 | L: 33 | L: 15 | L: 12 | Yes | Yes |
| Total: 36 | CL: 32, WID: 12, FID: 20 | L 883, R: 513 | L: 80, R: 28 | L: 64, R: 23 | L: 44, R: 41 | L: 7/10, R: 4/5 | L: 5/9, R: 3/5 |

Local field potentials were recorded via a GrapeVine Neural Interface (Ripple, LLC) sampling at 1 kHz. Notch filters at 60/120/180 Hz were applied online. Data was subsequently filtered from 0.1-115 Hz to isolate single trial potentials (stP) or decomposed via Morlet wave convolution to determine the power from 40-100 Hz to isolate single trial high frequency broad-band activity (stHFBB). These stHFBB responses were then Z-scored based on the baseline period from 500-0 ms proceeding stimulus onsets. It has been previously shown that these two aspects of the local-field potential, stP and stHFBB, contain complementary information (Miller et al., 2016), though also potentially arise from different neurophysiological generators (Engell et al., 2012; Hermes et al., 2012; Jacques et al., 2016; Leszczyński et al., 2020). Therefore, to assess the overall selectivity across VTC we use both as features in the classifiers described in *Multivariate temporal pattern analysis* (Figures 1B, 2-4, 6-8). We also investigated the independent contributions of these signal components to our category-selectivity maps (Figure 6). Trials where the stHFBB or stP exceeded 5 standard deviations from the mean were thought to contain noise and therefore excluded from further analysis.

**Figure 1.**
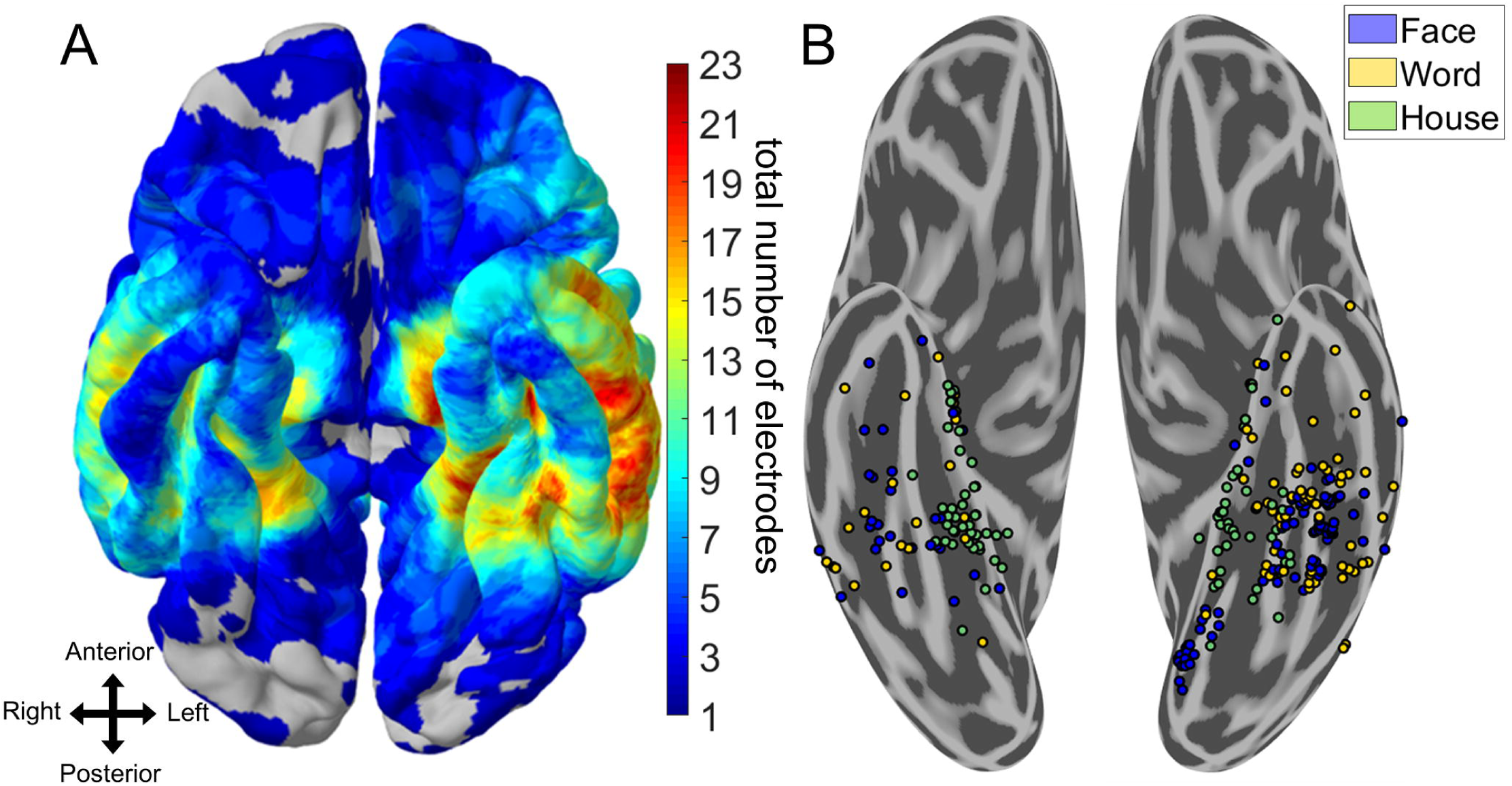
Spatial topography of word- and face-selective iEEG electrodes. A) Heat map of electrode coverage (both category-selective and non-selective) across 36 iEEG participants. Electrodes below the inferior temporal sulcus and anterior to the posterior edge of the fusiform gyrus on the participant’s native space were considered VTC. There was a greater proportion of left hemisphere coverage relative to right hemisphere coverage. B) Electrodes that responded preferentially to words, faces, or houses and could significantly discriminate these stimuli from all others using six-way Naïve Bayes classification (p < 0.05, Bonferroni corrected within participant). House-selective electrodes are primarily medial to word- and face-selective electrodes. Multiple adjoining word- and face-selective patches are found along the medio-lateral axis of ventral temporal cortex, bilaterally. Depth stereotactic EEG electrodes have been brought to the ventral surface for clarity.

#### Determining Language Laterality

Records from preclinical magnetoencephalography (MEG) language mapping sessions were used to determine the laterality of language function for 30 of the 36 iEEG participants. Language mapping records for the remainder of the participants could not be located. The preclinical language mapping records contained laboratory technician notes indicating whether MEG activity during reading, listening, and word-repetition tasks was lateralized to the left or right hemisphere. The original data from these sessions was not available to conduct more precise analyses of language laterality for these participants.

#### Multivariate temporal pattern analysis

To determine which electrodes contained information about word and face categories, leave-one trial out cross-validated Gaussian Naïve Bayes classifiers were used to predict the category of object participants were viewing given a sliding 100 ms of neural activity from one iEEG electrode during the category-localizer task (six-way classification). Signals from stP and stHFBB were both fed in as features to a single classifier for the main selectivity maps. This procedure was repeated from 100 ms prior to 900 ms after stimulus onset with 10 ms time-step to derive a time-course of decoding at each VTC electrode. We also ran separate classifiers on only features from stP or stHFBB to investigate the independent sources of information contained within these signal components. We ensured the number of features fed into these two types of classifiers was consistent by averaging 10 ms bins of stP, since stHFBB was sampled only every 10 ms, before classification.

Face-selective iEEG electrodes were defined as those that achieved a peak sensitivity (d’) of decoding for faces greater than the chance at the p < 0.05 level, Bonferroni corrected for multiple comparisons in time and across the total number of electrodes within a participant. Sensitivity (d’) describes the separation between a classifier’s noise and signal distributions and is defined as the inverse normal cumulative distribution function (Z’) of the true positive rate (TPR) minus the inverse normal cumulative distribution function of the false positive rate (FPR),

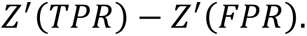

The Bonferroni corrected d’ sensitivity threshold was found by performing a binomial test on a null distribution of 1 million d’ sensitivities that were obtained by randomly classifying permutations of the trial labels. A small number of electrodes responded to all categories *except* faces, which resulted in above-chance face classification, since the distribution of responses to faces was significantly different from the responses to other object categories. Therefore, we imposed an additional criterion to determine selectivity: face-selective channels had to demonstrate a maximum peak event-related potential or event-related broadband response to faces relative to the other object categories. An identical procedure was done to define word- and house-selective electrodes.

To determine the independence of word- and face-selectivity within electrodes, we repeated the above multivariate pattern analysis for word- and face-selective electrodes after removing trials from the category they were most selective to. Word-selective electrodes were determined to also be selective for face stimuli if, after removing trials when words were presented, we could reliably predict trials where faces were presented from the other object categories (d’ sensitivity corresponding to p < 0.05, Bonferroni corrected for multiple temporal and electrode comparisons within participants using the same permutation test described above). Further, we stipulated that this d’ for faces must be greater than the d’ for all the remaining object categories. An identical procedure was used to define face-selective electrodes that were also selective for words.

To determine if word- and face-selective electrodes contained exemplar-level information about either faces or words, we performed pairwise classification of the face and word individuation stimuli for the electrodes on which we had data (Table 1). Specifically, in the case of word individuation, we used three-fold cross-validated Gaussian Naïve Bayes classifiers to predict which of two real words a participant was viewing based on sliding 100 ms of data from the word-selective electrodes. Three-fold cross-validation was used instead of leave-one-out cross validation (which was used for assessing category-level selectivity) to save computational time as there were many more models (stimulus pairs) tested with the exemplar classifier. We repeated this procedure across all pairs of real-words of the same length and averaged the time-courses of this pairwise decoding (56 pairs of words). We determined the p < 0.05 chance-level of this average pairwise decoding by repeating this procedure 1,000 times on data with shuffled trial labels in a subset of the word-selective electrodes (Maris and Oostenveld, 2007). These global null distributions were similar across the randomly subsampled electrodes and therefore we chose a d’ threshold corresponding to the highest p < 0.05 level obtained from this randomly chosen subset. We ran similar pairwise decoding and threshold definition on real-word versus pseudowords of the same length (36 pairs) and real-word versus false-font stimuli (136 pairs) to determine if electrodes that could not individuate real-words could perform these finer discriminations compared to those tested in the category localizer task.

Similarly, for face individuation we performed pairwise decoding of face stimuli during sliding 100 ms time-windows of face-selective electrode activity. We then averaged these time-courses across all 120 pairwise face classifications and calculated the p < 0.05 corrected level by repeating the permutation analysis described for the word individuation task on a random subset of face-selective electrodes.

#### Spatiotemporal k-means clustering

We used a spatiotemporal variant of k-means clustering to determine if spatially contiguous word- or face-selective regions demonstrated distinct temporal dynamics. For word- and face-selective electrodes, we separately standardized the d’ sensitivity time-courses derived from the category-level multivariate classifiers of left and right hemisphere electrodes from 100 to 600 ms post stimulus onset. We then concatenated this matrix with the electrodes’ MNI-coordinate, which was multiplied by a constant (spatial weighting parameter) that modulated the weight of the spatial versus temporal components of the signal to the clustering algorithm. We then performed k-means clustering using Euclidean distances and 100 repeats with random initializations to determine clusters of nearby word- or face-selective electrodes within each hemisphere that demonstrated correlated dynamics. Because the d’ time-courses were standardized, Euclidean distances were equivalent to correlation distance for the temporal data and Euclidean distance for the spatial data.

To determine the optimal weighting of spatial and temporal signal components and optimal number of clusters, we calculated the total spatial and temporal variance explained by the clustering solutions run with several spatial weighting parameters. This was performed for k = 1 to 10 clusters per hemisphere per faces or words. The elbow method was used to determine the optimal number of clusters per hemisphere. The optimal number of clusters was 4 for right hemisphere face-selective electrodes, 3 for right hemisphere word-selective electrodes, 3 for left hemisphere face-selective electrodes, and 4 for left hemisphere word-selective electrodes. We chose the spatial weighting parameter that explained the maximum amount of variance across k = 3 to 4 clusters per hemisphere per category (spatial weight = 300). Small deviations in the spatiotemporal weighting parameter did not strongly affect the overall organization of spatiotemporal clusters. The dynamics of these electrode clusters were then determined by averaging the selectivity time courses (d’ derived using *multivariate temporal pattern analysis*) across the electrodes belonging to each cluster.

#### Statistical analyses

Two sample t-tests were used to compare peak d’ sensitivity, peak latency, and onset latency for right versus left word- and face-selective electrodes. Onset latency was defined as the first time point that the d’ sensitivity reached a p < 0.001 threshold, which was non-parametrically defined using the d’ sensitivities of all object-selective electrodes from 500-0 ms prior to stimulus onset. Spearman’s rank-order correlations were used to test for relationships between peak d’ sensitivities and latency. We used linear mixed effects models to compare face and real word individuation in the category-selective clusters identified by the spatiotemporal k-means algorithm. Linear mixed effects models allowed us to determine if there were differences in peak individuation d’ or latency across these clusters while correcting for cross-subject differences. We only compared spatiotemporal clusters with greater than 10 electrodes with individuation data. The Satterthwaite approximation was used to estimate the degrees of freedom in these linear mixed effects models to compute the reported p-values. The time points corresponding to the leading edge of the classification window were used for all temporal statistical analyses.

### fMRI data collection and preprocessing

#### Participants

Eight participants (six females, mean age 25 years) participated in the fMRI experiment. All participants were right-handed, had normal or corrected to normal vision and gave written informed consent. The National institutes of Health Institutional review Board approved the consent and protocol (protocol 93 M-0170, clinical trials #NCT00001360). Participants were monetarily compensated for their time.

#### fMRI scanning parameters

All fMRI scans were conducted on a 7 T Siemens Mangetom scanner at the Clinical Research Center on the National Institutes of Health campus. Partial volumes of the occipital and temporal cortices were acquired using a 32-channel head-coil (42 slices, 1.2×1.2×1.2 mm; 10% interslice gap; TR=2s, TE=27ms; matrix size=170×170).

#### Experimental Paradigm

Participants fixated centrally whilst images of words, faces and houses were presented in blocks (16s per block). These images were taken from the same category localizer task presented to iEEG patients. In each block 20 exemplar stimuli were presented (300ms with a 500ms ISI). Participants performed a one-back task, responding, via MRI compatible response box, whenever the same image appeared twice in a row. Participants completed 10 runs of the localizer.

#### fMRI data preprocessing

All data were analyzed using the Analysis of Functional NeuroImages (AFNI) software package (Cox, 1996). Prior to statistical analysis, all images were motion corrected to the first volume of the first run. Post motion-correction data were detrended.

#### Statistical analysis

To identify word-, face- and house-selective regions, we performed a general linear model (GLM) analysis using the AFNI functions 3ddeconvolve and 3dREMLfit. The data at each time-point were treated as the sum of all effects thought to be present at that time point and the time series was compared against a Generalized Least Squares Regression model fit with REML estimation of the temporal auto-correlation structure. Responses were modelled by convolving a standard gamma function with a 16 second square wave for each condition (words, faces & houses). Estimated motion parameters were included as additional regressors of no-interest and fourth-order polynomials were included to account for any slow drifts in the MRI signal over time. Significance was determined by comparing the beta estimates for each condition (normalized by the grand mean of each voxel for each run) against baseline.

#### Split-half analysis

For each participant, the ten localizer runs were divided into odd and even splits. In each split, we performed the same GLM analysis as described above and looked for significant voxels for the contrast of words versus faces. Despite having only half of the data, we observed significant word-selectivity that was medial of face-selectivity consistently across participants. In order to quantify this selectivity in an independent manner, we first defined medial word-selective regions within a split (e.g. odd) and then sampled the data from the other half (e.g. even). ROIs were defined using data spatially smoothed with a 2 mm Gaussian kernel to generate spatially contiguous clusters, whereas the test data was not spatially smoothed. To avoid any bias in node selection, this process was then reversed and the average computed. Within each ROI we calculated the average t-value for each condition versus baseline.

## Results

From 1,396 intracranial electrode contacts implanted within or on the surface of VTC of 36 patients, we isolated those implanted in regions that were highly selective for either faces, words, or houses. Highly face-selective electrodes were defined as those that had both (1) single-trial responses that could significantly discriminate face presentations from presentations of five other object categories (words, houses, bodies, hammers, and phase-scrambled objects; p < 0.05 level, Bonferroni corrected for multiple spatial and temporal comparisons within participant; see *Materials and Methods*) and (2) responded maximally to faces compared to all other object categories on average. This ensured that electrodes designated as highly “face-selective” were those that responded maximally and were significantly selective for faces compared to the five other object categories. An identical procedure was used to define word- and house-selective electrodes.

108 electrodes demonstrated primarily face-selective responses (80 in the left, 28 in the right), 87 demonstrated primarily word-selective responses (64 in the left, 23 in the right), and 85 demonstrated primarily house-selective responses (44 in the left, and 41 in the right) (Figure 1). Figure 2 and Table 1 illustrate the distribution of object-selective electrodes across participants. The greater number of left versus right object-selective electrodes was comparable to the greater coverage of left VTC relative to right VTC in our patient population (883 electrodes implanted in the left, 513 in the right, Figure 1A). Although some word- and face-selective electrodes demonstrated partial selectivity for the other object category, there were several examples of electrodes that were strongly tuned to only words or faces (Figure 3). This suggests that the neural circuits responsible for processing words and faces are, at least, partially dissociable (Behrmann and Plaut, 2013; Susilo and Duchaine, 2013; Susilo et al., 2015).

**Figure 2.**
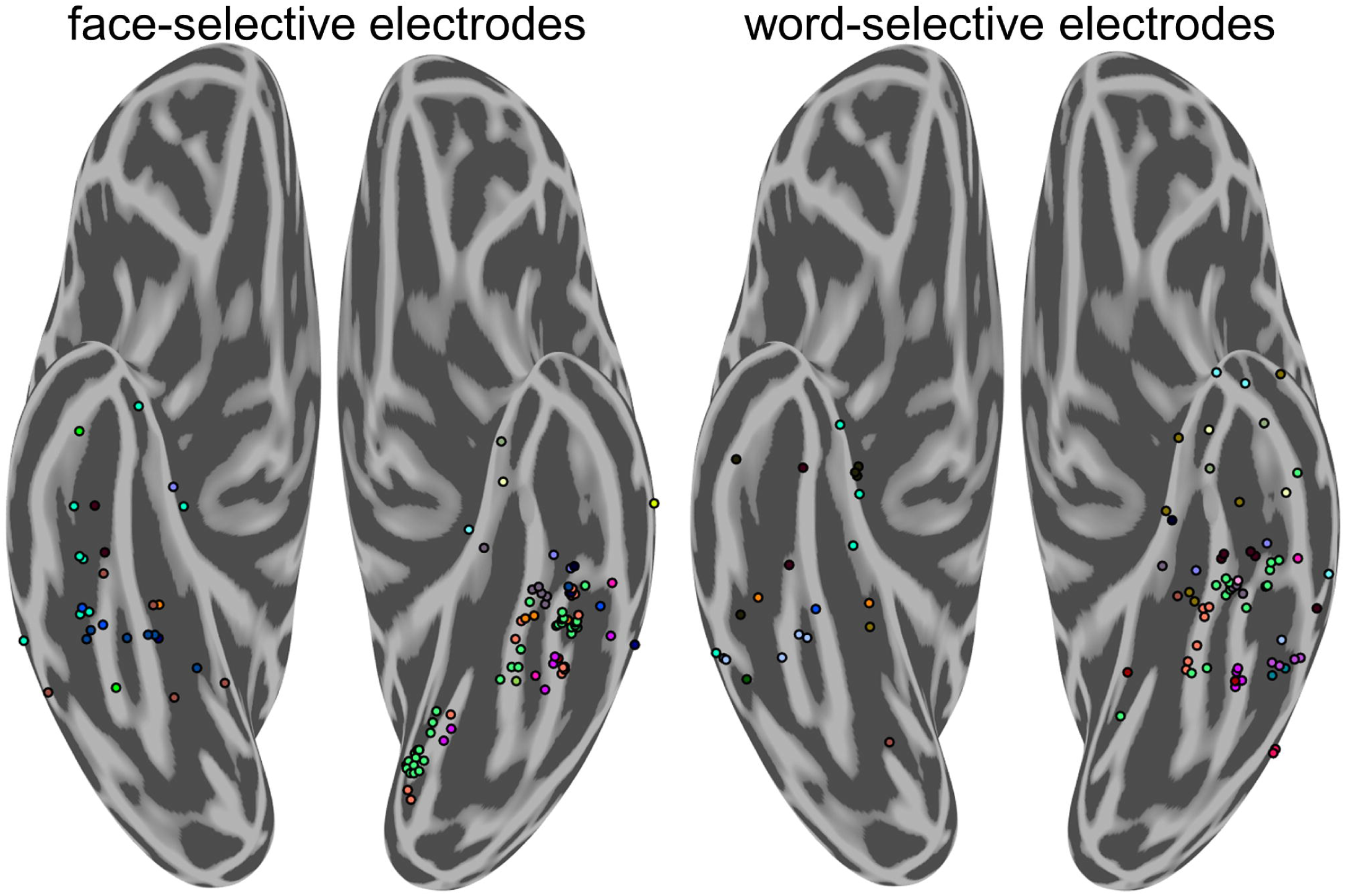
Distribution of face-selective and word-selective electrodes by participant. Distribution of highly face-selective (left) and word-selective (right) electrodes by participant. Color represents individual participants and corresponds across figure panels. Each group-level cluster of word- and face-selective electrodes is represented by data from multiple participants.

**Figure 3.**
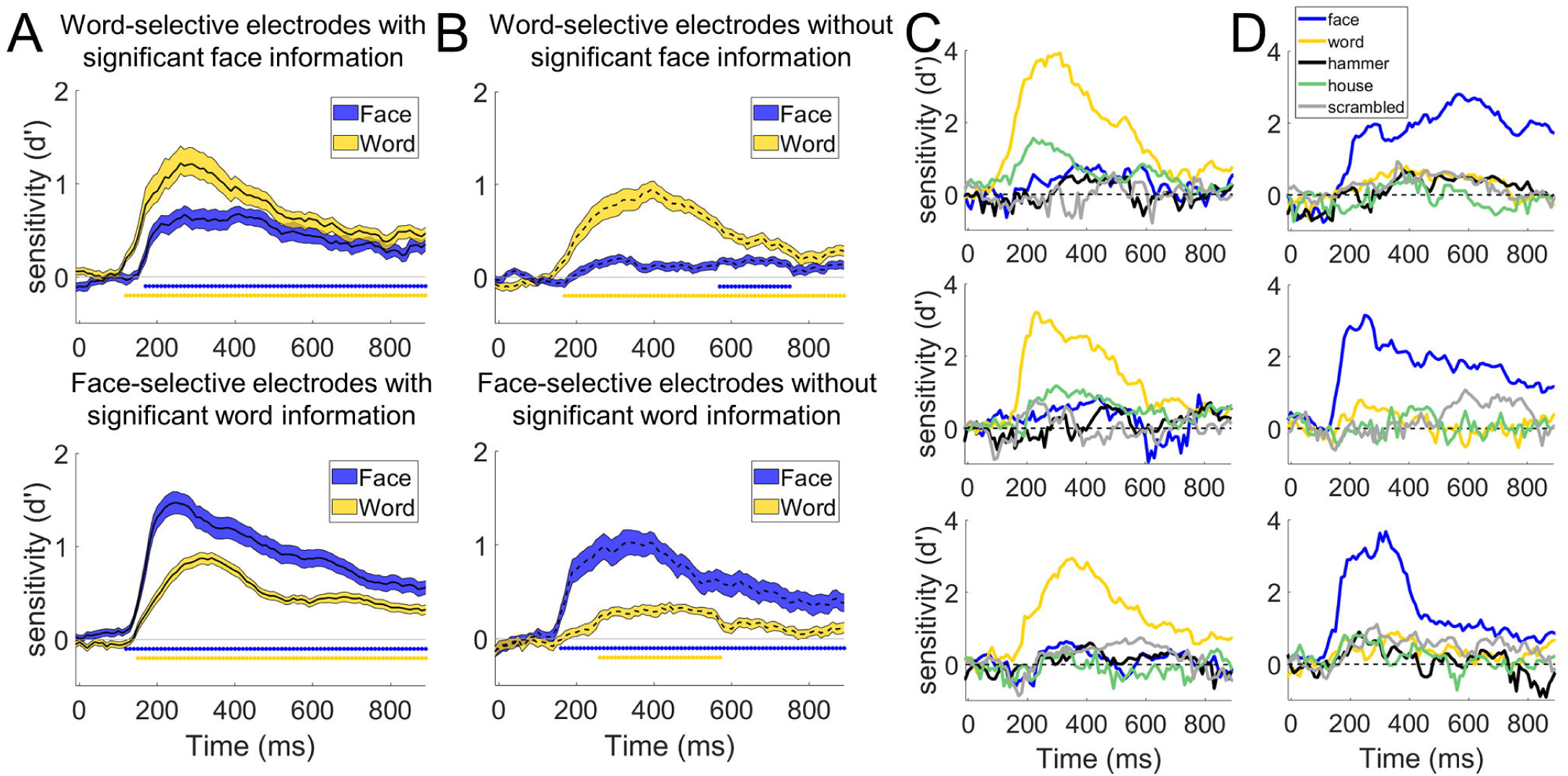
Independence of word- and face-processing networks. A) Average decoding time-course for word- (top, n = 39) and face- (bottom, n = 75) selective electrodes that contained significant amounts of information about the other object category. 21 out of 28 (75 %) face-selective electrodes in the right hemisphere and 54 out of 80 (66 %) in the left hemisphere could significantly discriminate words from the other object categories excluding faces (e.g. d’ sensitivity for words was above chance for 5-way classification of the non-face object categories) at the p < 0.05 level (Bonferroni corrected for multiple comparisons in time and electrodes within participant, see *Materials and Methods*). 9 out of 23 (39 %) word-selective electrodes in the right hemisphere and 30 out of 64 (47 %) in the left hemisphere could discriminate faces from the other object categories excluding words. Error bars indicate standard error from the mean across electrodes. Colored bars under the curves indicate times where the average selectivity is above chance (p < 0.001 corrected for temporal comparisons). B) Average decoding time-course for word- (top, n = 48) and face- (bottom, n = 33) selective electrodes that did not contain above chance information for the other object category. Although decoding accuracy was above chance at later time points for the non-preferred category across the population of electrodes, decoding accuracy was much smaller for the non-preferred compared to preferred category. C) Example decoding time courses from three highly word-selective electrodes that did not display face-selectivity. D) Decoding time courses of three highly face-selective electrodes that did not display word-selectivity. The patient from which the middle electrode recording was obtained was not presented with pictures of hammers.

To assess how word- and face-processing networks organize relative to one another, the spatial topography of word-, face-, and house-selective electrodes was examined. At the group level, selectivity to house stimuli was found primarily along the left and right parahippocampal gyrus, with some cases where selectivity extended into the collateral sulcus and medial fusiform gyrus. These patches were generally medial to word- and face-selective locations, consistent with previous fMRI and iEEG studies (Halgren et al., 1994; Haxby et al., 1996; Aguirre et al., 1998; Cohen et al., 2000; Kadipasaoglu et al., 2016). Face-selectivity was found primarily along the left and right fusiform gyrus with some face-selective regions within the lingual gyrus, and occipitotemporal sulcus (Figure 1B). Consistent with prior findings (Cohen et al., 2000), word-selective regions were found on the lateral bank of the fusiform and into the occipitotemporal sulcus in the left hemisphere. Word-selective regions were also found anterior to most prior reports from fMRI, in locations that generally have poor signal due to susceptibility artifacts (Devlin et al., 2000). In contrast to most maps of word- and face-selective regions obtained from fMRI (Allison et al., 1994; Haxby et al., 1994; Puce et al., 1996; Harris et al., 2016; Saygin et al., 2016; Dehaene-Lambertz et al., 2018; Gomez et al., 2018), a mosaic of word-selective regions were also found medial to face-selective regions, on the medial bank of the fusiform and into the collateral sulcus. Each of these face-, word-, and house-selective regions were found in multiple participants (Figure 2), demonstrating relatively consistent localization of these regions at a group level.

Interdigitation of word- and face-selective regions was seen in the left hemisphere of 5 out of 9 participants with at least two word-selective electrodes and one face-selective electrode or vice-versa and in the right hemisphere of 3 out of 5 such participants (Table 1, see Figure 4 for examples). Word-selective regions were found strictly medial to face-selective regions in the left hemisphere of 7 out of 10 participants with at least one word- and one face-selective electrode and in right hemisphere of 4 out of 5 participants (Table 1, see Figure 4 for an example). Thus, highly word-selective regions medial to face-selective regions was not simply a consequence of individual variability in a group-level map but instead was detected in the majority of participants that had coverage of both face- and word-selective VTC.

**Figure 4.**
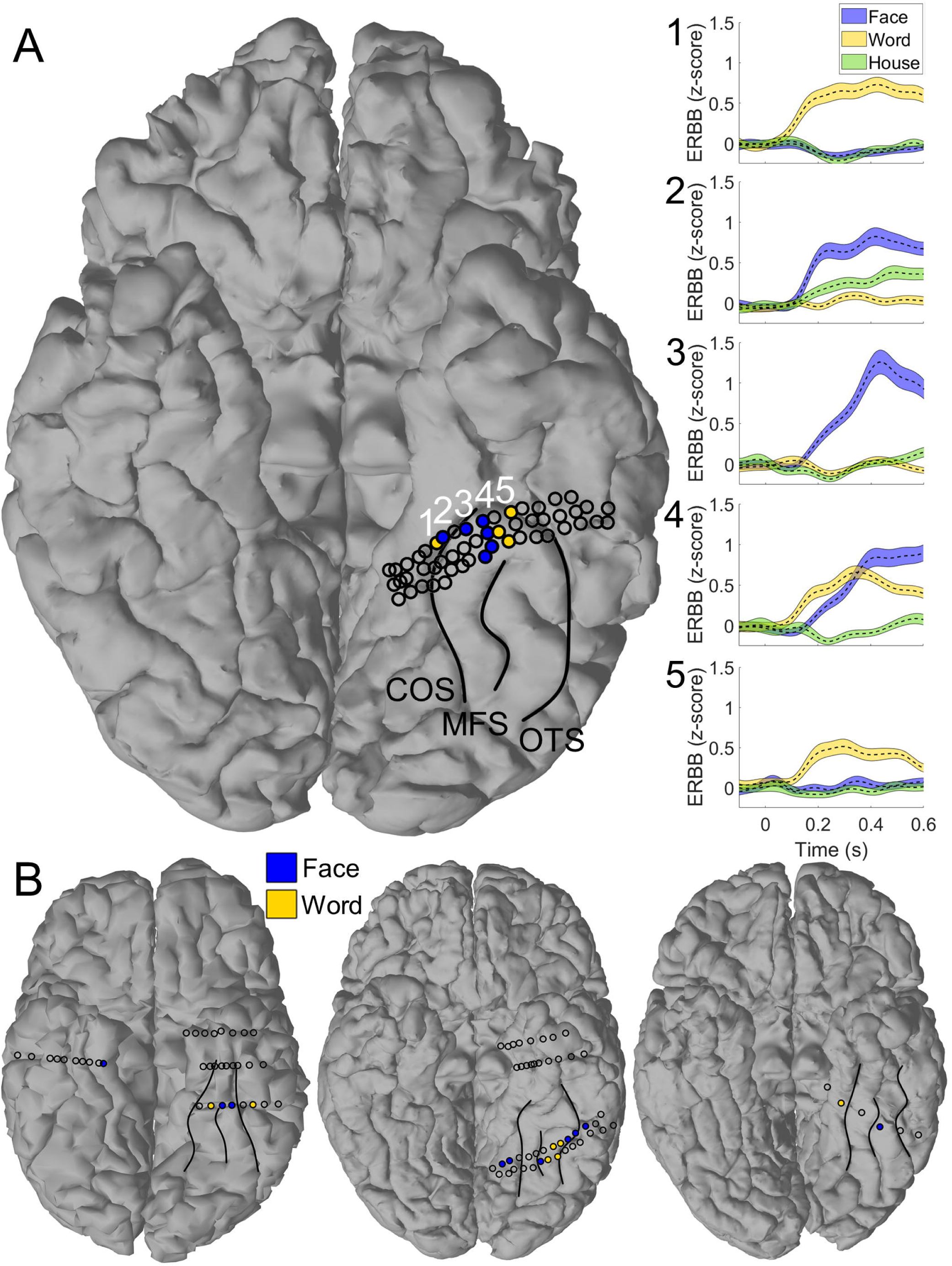
Multiple adjoining word- and face-selective patches in individual participants. A) Representative single participant demonstrated alternating bands of word- and face-selectivity along the left fusiform gyrus. Shaded electrodes are those selective to words (yellow) and faces (blue). Non-filled circles represent ventral temporal electrodes that did not reach the selectivity criterion for either of these categories. Raw event-related broadband activity is shown for each of the numbered electrodes in the right panel. Moving from medial to lateral, electrodes demonstrate a preferential response to words, mixed response to both words and faces, preferential response to faces then preferential response to words. Abbreviations: collateral sulcus (COS), midfusiform sulcus (MFS), occipitotemporal sulcus (OTS). B) Three additional examples of patients with multiple adjoining word- and face-selective regions or word-selectivity medial to face-selectivity in VTC. Major VTC sulci have been labeled for clarity.

Because word-selective patches were found medial to face-selective patches in the iEEG data, which is generally not observed in 3 T fMRI studies (Haxby et al., 1994; Puce et al., 1996; Dehaene et al., 2010), we sought to determine if a similar organization existed in healthy participants using the higher resolution of 7 T fMRI. When contrasting responses to words and faces in eight participants, face-selectivity was primarily centered on the midfusiform sulcus while word-selectivity was greatest in the occipitotemporal sulcus (Figure 5). Consistent with the iEEG results, six of the eight participants demonstrated left word-selective regions medial to face-selective regions on the fusiform gyrus. In these medial word-selective patches, responses to words were significantly greater than responses to both face and house stimuli (p < 0.001, split-halves analysis). These medial word-selective regions were approximately 1/3^rd^ the size of more lateral word-selective regions (mean size of lateral word-selective regions: 398 voxels; std. error: 43 versus medial regions: 139 voxels; std. error: 29 voxels; p < 0.01). Also, 7 out of 8 of the healthy participants demonstrated word-selective patches near the anterior tip of the fusiform, despite susceptibility artifacts (Devlin et al., 2000), consistent with the iEEG data (Figure 1B). Altogether, the map of word- and face-selective regions of the left hemisphere derived from 7 T fMRI were consistent with those derived from iEEG, medial and anterior word-selective regions are not seen in most maps drawn from 3 T fMRI (Haxby et al., 1994; Puce et al., 1996; Dehaene et al., 2010).

**Figure 5.**
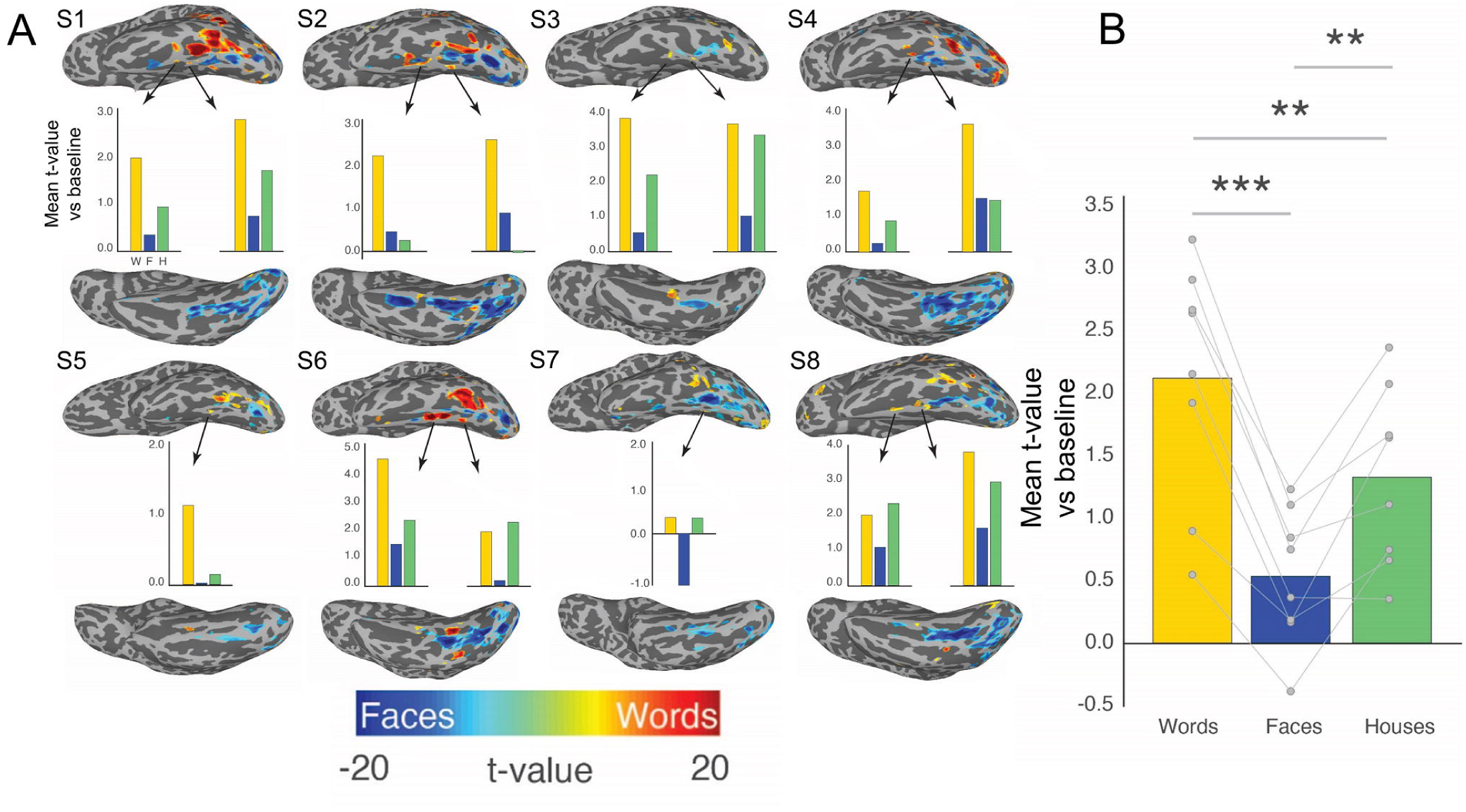
Interdigitation of BOLD responses to words and faces across eight healthy participants. Eight healthy participants that underwent an identical category localizer task as the iEEG participants demonstrated similar category selectivity. A) Individual maps demonstrate word versus face-selectivity across left VTC. In six out of eight of these participants there was strong word-selectivity medial to face-selectivity along the midfusiform sulcus. The bar graphs below each participant indicates the selectivity of these word-selective regions when defining them based on one half of the data and testing on the other half of the data. Word-selective responses were less consistent in the right hemisphere across participants. B) Bar graph of word-selectivity in left hemisphere medial word-selective regions across participants for the left-out half of the data. Symbols: ** p < 0.01, *** p < 0.001.

The maps in Figures 1-3 were made by combining two key aspects of the iEEG signals, the single trial potentials (stP) and the single trial high frequency broadband activity (stHFBB), to examine the category-selectivity of the underlying VTC neural populations in aggregate across these signal components. Studies have shown that while category-selectivity demonstrated in stP and stHFBB often overlaps, they are not redundant (Engell and McCarthy, 2011; Engell et al., 2012; Miller et al., 2016), suggesting that stP and stHFBB have at least partially distinct physiological generators. To examine these signal components separately, we trained multivariate classifiers solely on stP or stHFBB and isolated electrodes that were selective in either signal component using the same criteria as before (single-trial discriminability and highest signal amplitude for words, faces, or houses). 58 electrodes showed significant selectivity in both stP and stHFBB (Figure 6A). Notably, the regions that demonstrated selectivity in both stP and stHFBB were those most often identified in canonical maps of category-selectivity based on fMRI (Cohen et al., 2000; Vinckier et al., 2007; Tsao et al., 2008; Weiner and Grill-Spector, 2010; Lerma-Usabiaga et al., 2018). Specifically, house-selectivity was restricted to the parahippocampal cortex, face-selectivity was primarily restricted to the fusiform bilaterally, and word-selectivity was restricted primarily to the left posterior-lateral fusiform and occipitotemporal sulcus. Regions that were less consistent with canonical fMRI maps tended to be those that were not significantly selective in both stP and stHFBB. For example, the medial word-selective patches were primarily seen in stP alone (Figure 6B), whereas anterior and right hemisphere word-selectivity was prevalent in either stP or stHFBB alone (Figures 6B and 6C). Broadly, more electrodes demonstrated selectivity in stP (232 electrodes from 32 participants, Figure 6B) compared to stHFBB (115 electrodes from 24 participants, Figure 6C). More widespread stP selectivity is consistent with a previous study comparing stP and stHFBB responses for faces in VTC, though that study did not observe any cases where selectivity for faces was demonstrated in stHFBB but not stP (Engell and McCarthy, 2011). The similarities and differences in selectivity demonstrated in stHFBB and stP are consistent with the hypothesis that these signals have different physiological generators (Lachaux et al., 2005), which may differ in their laminar distribution (Leszczyński et al., 2020) and spatial signal-to-noise falloff (Engell and McCarthy, 2011). Additionally, different category-selectivity across these iEEG signal components may also help explain differences between category-selectivity maps drawn from iEEG and fMRI, as some studies suggest fMRI has differential sensitivity to these aspects of the iEEG signal (Conner et al., 2011; Engell et al., 2012; Jacques et al., 2016).

**Figure 6.**
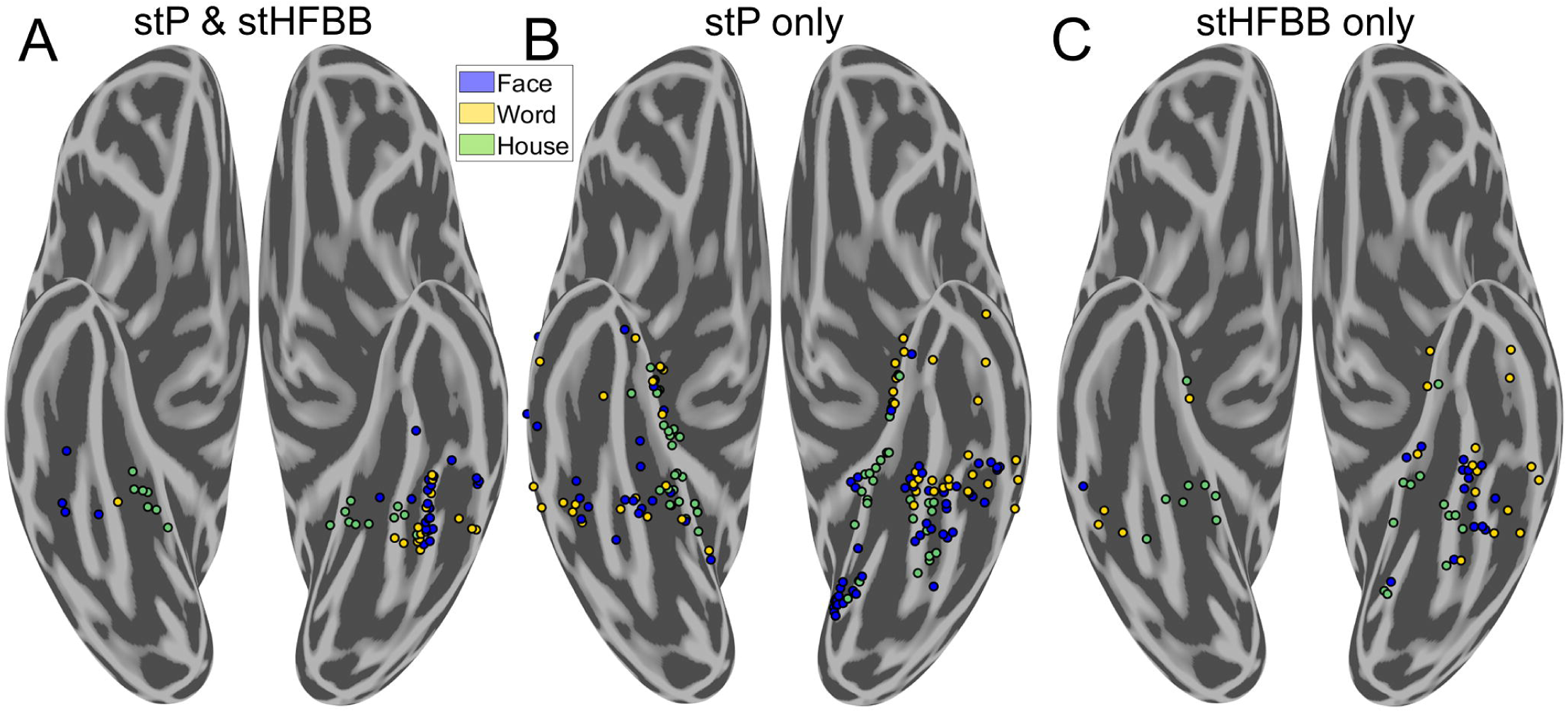
Comparing category-selectivity in single-trial potentials and high-frequency broadband. Differing spatial distribution of electrodes that demonstrated selectivity in single-trial potentials (stP) and single-trial high-frequency broadband activity (stHFBB). A) Electrodes that demonstrated selectivity in both stP and stHFBB were clustered around the fusiform and parahippocampal gyri. B) Electrodes selective in only stP were much more widely distributed and included medial and anterior word-selective regions not typically seen in fMRI. C) Electrodes that were only selective in stHFBB were less prevalent than those only selective in stP, but also extended outside of the fusiform and parahippocampal gyri.

One question is whether word- and face-selective regions identified using iEEG discriminate between individual face and word exemplars, respectively. Classifying at the exemplar level also can address the potential concern that the word- and face-selective regions identified using iEEG may be responding to low-level features that drastically differ between the sampled image categories. A subset of the iEEG participants underwent independent word and face individuation tasks (see *Materials and Methods,* Table 1). Activity from 85 out of 97 sampled face-selective electrodes in 13 participants could be used to reliably predict the identity of a presented face. Similarly, activity from 40 out of 53 sampled word-selective electrodes from 10 participants could be used to discriminate single words of the same length from one another. Of those 13 word-selective electrodes that could not reliably achieve word individuation, six could reliably discriminate pseudowords from real words of the same length, seven could reliably discriminate false-fonts from real words. Therefore, most of the word- and face-selective regions mapped with iEEG contained reliable exemplar-level information specific to the categories they were selective to.

Peak word and face individuation was significantly correlated with peak category-selectivity in word and face-selective regions for which we had individuation data (word-selective: Spearman’s ρ(53) = 0.50, p < 0.0001, face-selective: ρ(97) = 0.48, p < 0.0001). Note that correlations in peak category-selectivity and within-category individuation may arise due to similar differences in measurement noise across recording contacts (for example, due to the distance the electrode was placed from the underlying face or word selective neural groups), underlying neural/physiological factors, or some mix of both.

In addition to the medial band of word-selective regions, there were a high proportion of right word-selective electrodes in our iEEG population (Figure 1B, Table 1). Although this finding is consistent with some other fMRI (Ben-Shachar et al., 2007; White et al., 2019) and iEEG studies (Halgren et al., 1994; Lochy et al., 2018), right hemisphere word-selectivity is often not seen in neuroimaging (Cohen et al., 2000, 2002) and was not very strong in our 7 T fMRI data either (Figure 5). 23 word-selective electrodes were found across nine participants in right VTC, out of 21 participants with right VTC object-selectivity. This discrepancy between right word-selectivity observed in fMRI and iEEG was also not attributable to participant handedness, since no participant with right word-selective regions was lefthanded. Three out of nine of these participants demonstrated evidence for bilateral language function while the other six demonstrated left dominant language function determined by preclinical magnetoencephalography (MEG, see *Materials and Methods*). Across the entire participant population, seven out of 30 iEEG participants with preclinical MEG demonstrated bilateral language function, the others were considered left dominant. One participant with bilateral language function and right hemisphere object-selectivity did not demonstrate right word-selectivity. Overall, neither participant handedness nor language dominance sufficiently explains the high proportion of word-selective regions found in right VTC.

While neither language laterality nor handedness explained right word-selectivity, substantial differences were seen in the dynamics of neural activity recorded from left versus right word-selective regions (Figure 7). Latency to word-selectivity onset and peak was shorter in left compared to right hemisphere word-selective regions (mean onset latency difference +/- 95 % CI: −133 +/- 61 ms, T(85) = −4.4, p < 0.0001, mean peak latency difference: −138 +/- 63 ms, T(85) = −4.3, p < 0.0001, Figure 7). These relationships held when taking into account potential differences in posterior to anterior coordinate of word-selective regions across hemispheres (onset: T(85) = −4.01, p = 0.0001, peak: T(85) = −3.97, p = 0.0002). There was no significant difference between the latency to peak d’ sensitivity or sensitivity onset for right and left face-selective regions (mean onset latency difference: −29 +/- 53 ms, T(106) = −1.1, p = 0.28, mean peak latency difference: 18 +/- 57 ms, T(106) = 0.63, p = 0.53, Figure 7). Additionally, the amplitude of peak d’ sensitivity for words was significantly greater in the left compared to right hemisphere word-selective regions (mean peak d’ sensitivity difference: 0.66 +/- 0.37, T(85) = 3.5, p = 0.0006). The amplitude of peak d’ sensitivity to faces was also significantly greater in the left compared to right hemisphere face-selective regions (mean peak d’ sensitivity difference: 0.58 +/- 0.39, T(85) = 3.0, p = 0.0037). There was a significant correlation between peak latency and peak magnitude within face-selective regions in the left (ρ(80) = −0.61, p < 0.0001) and right (ρ(28) = −0.79, p < 0.0001) hemisphere and word-selective regions in the left (ρ(64) = −0.68, p < 0.0001), but not right (ρ(23) = −0.15, p = 0.48) hemisphere, suggesting that longer peak latencies were associated with smaller peak selectivity. These correlations were not significantly different between face-selective regions in the left and right hemisphere (T(85) = −1.56, p = 0.058), but there was a greater correlation between peak latency and magnitude in left compared to right hemisphere word-selective regions (T(85) = 2.63, p = 0.004) Given that it was only true for word-selective electrodes, the relatively slower response of right versus left word-selective regions may potentially explain differences in word-selectivity maps derived from iEEG and fMRI and may reflect the left hemisphere bias for language.

**Figure 7.**
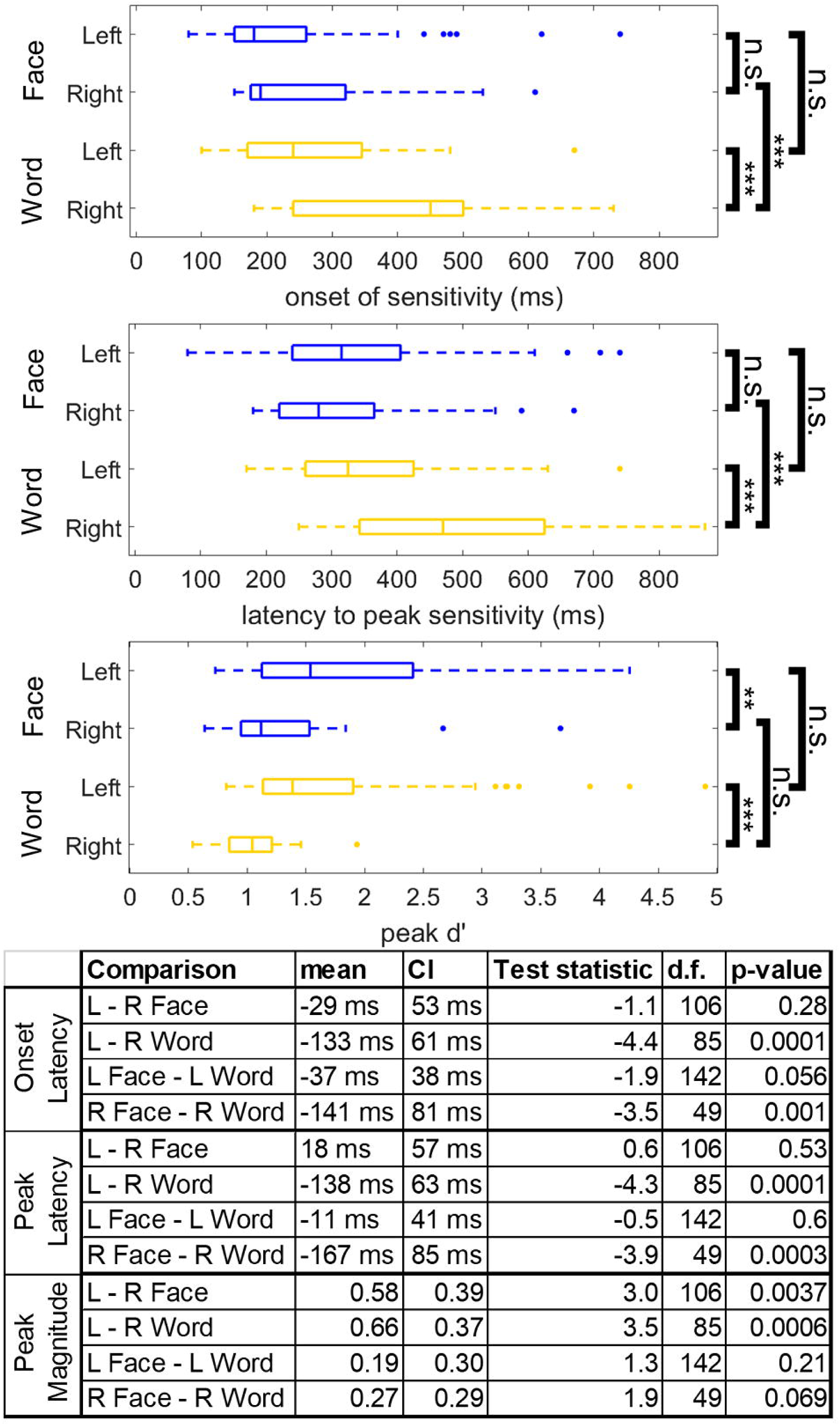
Differences in the dynamics of left versus right word- and face-selective regions. Latency of word (yellow) and face (blue) sensitivity onset, latency of peak sensitivity, and magnitude of peak sensitivity across hemispheres. Latency of sensitivity onset is defined as the first timepoint that reached a d’ corresponding to p < 0.001 non-parametrically defined by the pre-stimulus baseline period. All time points reference the leading edge of the classification window. Box plots represent median, 25^th^ and 75^th^ percentiles. Summary statistics of each box plot are presented in the table. Abbreviations: confidence interval (CI), degrees of freedom (d.f.), left (L), right (R). Symbols: n.s. p > 0.05, * p < 0.05, ** p < 0.01, *** p < 0.001.

Finally, using the iEEG data, we sought to determine if there were any differences in the temporal dynamics of neural responses across word or face-selective regions within the same hemisphere. We used a spatiotemporal k-means clustering algorithm to find spatially contiguous regions of left and right VTC which demonstrated correlated category-selective dynamics. After optimizing the algorithm to capture the most spatiotemporal variance with the optimal number of clusters (see *Materials and Methods*), we could compare the dynamics of distinct word- and face-selective clusters within VTC.

Word-selective regions were clustered into 4 distinct left hemisphere clusters and 3 right hemisphere clusters (Figure 8A). Word-selective regions on the left fusiform gyrus demonstrated the earliest and strongest selectivity, peaking around 200 ms (Figure 8B, gray). Left hemisphere medial word-selective regions and right hemisphere word-selective regions came next, peaking around 300 ms (Figure 8B, green and cyan) followed by lateral regions around 350 ms (Figure 8B, red). Word-selective regions in left anterior VTC peaked around 400-450 ms (Figure 8B, blue); right more anterior regions peaked around 600 ms (Figure 8B, magenta). When considering word-selectivity dynamics exhibited independently in stP and stHFBB signal components, word-selective electrodes on the fusiform demonstrated strong selectivity in both signal components, whereas other regions displayed distinct dynamics across these signal components (Figures 7C-D).

**Figure 8.**
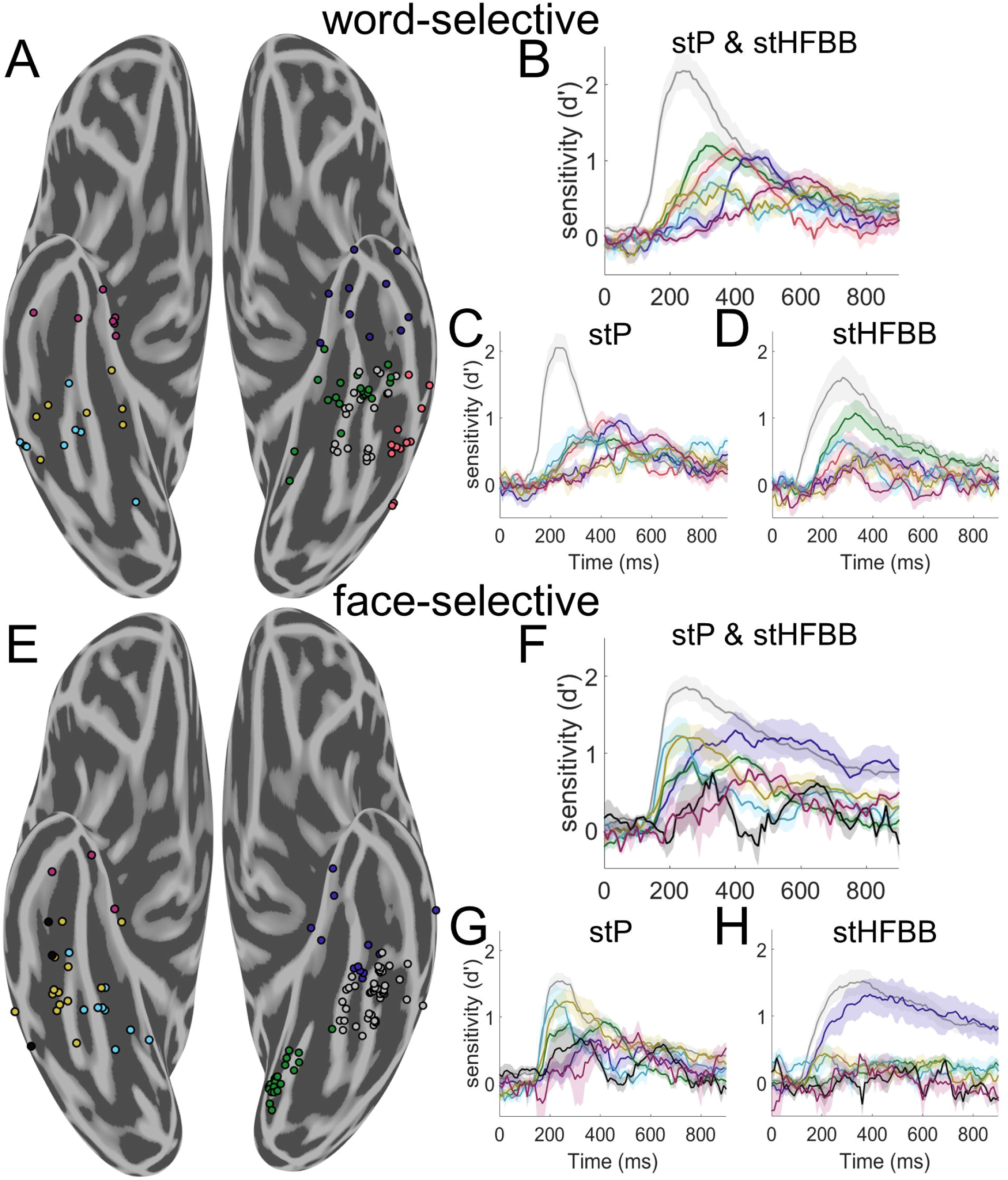
Spatiotemporal clustering of word- and face-selective regions. A) Spatiotemporal clustering of word-selective VTC electrodes. The illustrated clustering solution was robust to different weightings of spatial and temporal information. Left hemisphere word-selective electrodes were clustered into four spatial clusters. A cluster was found on the fusiform gyrus (gray, 21 electrodes from 5 participants), as well as medial (green, 20 electrodes from 10 participants), anterior (blue, 11 electrodes from 5 participants), and lateral (red, 12 electrodes from 7 participants) to the fusiform gyrus. Right hemisphere word-selective regions had later onsets and were more clearly separated along the posterior to anterior axis (posterior: cyan; 8 electrodes from 4 participants, mid: yellow; 8 electrodes from 3 participants, anterior: magenta; 7 electrodes from 6 participants). B) Average d’ timecourse of each group of electrodes in A when jointly classifying stP and stHFBB. Error bars represent standard error across electrodes. C) Average d’ time course of each group of electrodes when classifying only stP. D) Average d’ time course of each electrodes when classifying only stBB. Word-selective electrodes on the fusiform demonstrate strong selectivity in both stP and stHFBB, whereas other regions display distinct dynamics across these signal components. E) Spatiotemporal clustering of face-selective VTC electrodes. Left hemisphere electrodes were clustered into three spatial clusters roughly posterior to (green, 21 electrodes from 3 participants), on (gray, 46 electrodes from 12 participants), and anterior to the fusiform gyrus (blue, 13 electrodes from 7 participants). Right hemisphere, face-selective electrodes were primarily clustered along the posterior to anterior VTC axis into four clusters (posterior: cyan; 9 electrodes from 5 participants, mid: yellow; 13 electrodes from 6 participants and black; 3 electrodes from 2 participants, anterior: magenta; n = 3 electrodes from 3 participants). F) Average d’ time course of each group of electrodes illustrated in E when jointly classifying stP and stHFBB. G) Average d’ timecourse of each group of electrodes when classifying only stP. H) Average d’ time course of each group of electrodes when classifying only stBB. Face-selective electrodes on the fusiform demonstrate strong selectivity in both stP and stHFBB, whereas other regions display distinct dynamics across these signal components.

Face-selective regions were organized into 3 distinct clusters in the left hemisphere and 4 distinct clusters in the right hemisphere (Figure 8E). Face-selective regions of the left and right fusiform gyrus demonstrated the earliest and largest peak selectivity around 200-250 ms (Figure 8F, gray and cyan). More anterior right hemisphere regions and a cluster of electrodes in left posteromedial VTC (Figure 8F, yellow and green) peaked around 300 ms. Finally, more anterior face-selective electrodes in left and right VTC peaked around 400 ms (Figure 8F, blue, black, and magenta). When considering face-selectivity dynamics exhibited independently in stP and stHFBB signal components, electrodes on the fusiform demonstrated strong selectivity in both components, whereas other regions displayed distinct dynamics across these signal components (Figures 7G-H).

From electrodes sampled in the word individuation task, we observed stronger word individuation in left word-selective regions on the fusiform compared to the more medial word-selective cluster illustrated in Figure 8A (peak d’ of fusiform minus medial regions: T(30) = 3.62, p = 0.001, linear mixed-effects model). There was no significant difference between the latency to peak word individuation across these clusters (T(30) = 2.91, p = 0.68). There were not sufficient subjects with electrodes in the other word-selective clusters with word individuation data to make comparisons between all clusters. Neither peak face individuation (T(50) = 1.03, p = 0.31) nor latency to peak face individuation (T(50) = −0.21, p = 0.84) was significantly different between face-selective regions along the left fusiform gyrus and the posteromedial face-selective cluster observed in Figure 8E. There were not sufficient subjects with electrodes in the other face-selective clusters with face individuation data to make comparisons between all clusters.

Overall, for both faces and words, these results suggest a cascade of processing that begins in the fusiform. Notably, the dynamics of these clusters suggest that they contribute to distinct stages of face- and word-processing, since the latencies of their responses are far longer than would be expected from feedforward visual transmission delays alone (Thorpe et al., 1996; Kravitz et al., 2013), but not long enough to exclude them from being relevant to perceptual behavior (Quian Quiroga et al., 2008; Tang et al., 2014).

## Discussion

In the current study, we found several VTC regions that demonstrated strong word-, face- and house-selective responses. Although activity recorded from VTC electrodes often contained information about multiple object categories, several selectively responded only to faces or words (Figure 3). Electrodes which demonstrated preference to only words or faces suggests that VTC word- and face-processing networks are not entirely overlapping (Behrmann and Plaut, 2013), but instead involve at least some independent nodes (Susilo and Duchaine, 2013; Susilo et al., 2015), which is also supported by stimulation and lesion evidence (Hirshorn et al. 2016).

In both the iEEG and fMRI data, strong face-selectivity along the fusiform gyrus was adjoining with highly word-selective regions in and around the occipitotemporal and collateral sulci. House-selective regions were found primarily along the parahippocampal gyrus. This organization of house-versus word- and face-selective regions supports that typical retinotopic eccentricity is an important organizing principle of VTC (Grill-Spector and Weiner, 2014). The word-selective regions around the occipitotemporal sulcus are consistent with prior studies showing word-selectivity within lateral aspects of (Dehaene et al., 2002; Price and Devlin, 2003). Due to sparse and variable sampling across participants, the data cannot address the question of whether there is a gradient of word-selectivity along the occipitotemporal sulcus (Vinckier et al., 2007) or distinct patches (Lerma-Usabiaga et al., 2018; White et al., 2019).

Despite some similarities with previous neuroimaging work, the iEEG and 7 T fMRI data here are inconsistent with a map of VTC wherein word-selective regions are strictly lateral to face-selective regions (Haxby et al., 1994; Puce et al., 1996; Dehaene et al., 2010). While there has been some mixed reporting of word-selectivity in anterior and medial VTC regions (Allison et al., 1994; Haxby et al., 1994; Puce et al., 1996; Harris et al., 2016; Saygin et al., 2016; Dehaene-Lambertz et al., 2018; Gomez et al., 2018), most models of orthographic-processing within VTC consider only the more lateral, traditional “visual word form area” (Dehaene et al., 2002; Price and Devlin, 2003). The disagreement between the observed organization of face- and word-processing networks in VTC and most previous maps drawn from fMRI may be the product of spatial smoothing commonly applied during fMRI data analysis (Geissler et al., 2005), signal dropout induced by susceptibility artifacts (Devlin et al., 2000), or the inferior sensitivity of 3 T fMRI relative to 7 T fMRI. Here, a mosaic of word-selective regions was found medial and anterior to face-selective regions at the group level and within multiple iEEG patients and in 7 T fMRI in healthy individuals. This evidence makes it unlikely that our observations are the product of inter-participant variability or differences between healthy controls and patients with intractable epilepsy (see also (Matsuo et al., 2015; Jonas et al., 2016; Kadipasaoglu et al., 2016; Lochy et al., 2018)). This mosaic organization of visual word-selective regions is similar to the mosaic organization of auditory language processing networks (Flinker et al., 2011), suggesting this pattern of organization may not be specific to the visual system.

Medial word-selective regions may reflect differential mediolateral VTC selectivity to object rectilinearity, with more medial VTC being more responsive to straight over curvy objects (Srihasam et al., 2014; Bao et al., 2020). However, the interdigitation of word- and face-selective regions along the mediolateral axis is not well captured solely by this rectilinear model or the retinotopic model. Instead, medial and lateral word-selective regions with distinct dynamics may indicate an interaction between multiple representational axes in VTC (Konkle and Caramazza, 2013; Grill-Spector and Weiner, 2014) and possibly competition between faces and words for cortical space (Behrmann and Plaut, 2020).

Previous studies have used electrical stimulation to demonstrate that a large portion of VTC, sometimes termed the “basal temporal language area,” plays a role in language processing (Krauss et al., 1996; Mani et al., 2008; Fonseca et al., 2009; Enatsu et al., 2017). However, generalized language deficits after lesions of the basal temporal language area are relatively minor (Krauss et al., 1996) and the relationship between reading deficits and VTC lesions, other than the visual word form area (Gaillard et al., 2006; Hirshorn et al., 2016), is unclear. A recent study reported differential language-related deficits during reading, repetition, and picture naming depending on the area of VTC stimulated (Forseth et al., 2018). Future studies are necessary to understand the precise relationship between medial, lateral, and anterior word-selective VTC dynamics and these regions’ functional contribution to reading and/or language processing.

Category-selective regions most consistent with prior fMRI studies were those that demonstrated selectivity in both stHFBB and stP iEEG signal components. In contrast, we found that medial word-selectivity was primarily demonstrated in stP rather than stHFBB. Previous studies have suggested that fMRI BOLD have differential sensitivity to stHFBB versus stP (Hermes et al., 2012), with some suggesting greater sensitivity to stHFBB (Engell et al., 2012; Jacques et al., 2016). Differential sensitivity to stP and stHFBB may explain why previous fMRI studies have only inconsistently observed medial word-selective regions. Our 7 T fMRI data shows that, with adequate power, both lateral and medial word-selective regions are seen in the left hemisphere using BOLD within individual participants. Future studies are necessary to fully understand the functional characteristics and neurophysiological generators of stP and stHFBB iEEG components (Miller, 2010; Ray and Maunsell, 2011; Leszczyński et al., 2020) and how they relate to any differential roles that medial and lateral word-selective regions play in reading.

In addition to this complex organization of word- and face-selectivity within hemispheres, our iEEG analyses suggest that right word-selective regions demonstrate longer latencies and lower amplitudes of peak selectivity compared to left word-selective regions, which may reflect the primary role the left, language dominant, hemisphere plays in word-processing (Fiez and Petersen, 1998). Previous studies have demonstrated weaker correlations between object-selectivity measured with iEEG and fMRI correlations at later time windows (Jacques et al., 2016). This may explain why bilateral selectivity to words is inconsistent across neuroimaging studies.

It has previously been suggested that right word-selective regions (along with left posterior word-selective regions) are involved in relatively early visual processing of words and then this information flows to left anterior word-selective regions (White et al., 2019). However, the dynamics observed here do not support this hypothesis, because left word-selectivity substantially preceded right word-selectivity. Instead, the time-course of right hemisphere activation is coincident with P300 and N400 potentials observed during reading, suggesting that right hemisphere word-selective regions may support the left hemisphere in later computations, such as those involving word syntax, memory encoding, and/or semantic processing (Friedman et al., 1975; Kutas and Hillyard, 1980; Federmeier and Kutas, 1999; Otten and Donchin, 2000; Arbel et al., 2011).

Word- and face-selective regions within hemispheres also demonstrated distinct dynamics. Word-selective regions on the left fusiform gyrus demonstrated the earliest and strongest word-selective responses. This was followed by word-selective activity in left occipitotemporal and collateral sulcus as well as right posterior word-selective regions. Finally, word-selective activity spread to anterior VTC between 400-600 ms. Further, the relatively later responses of word-selective regions outside of the fusiform may also contribute to differences in category-selective maps drawn from iEEG and fMRI (Jacques et al., 2016).

Face-selective responses were strongest and earliest on the fusiform gyrus bilaterally. A cluster of posteromedial face-selective electrodes was found in early visual cortex. The slower time-course of these regions compared to face-selective regions on the fusiform suggests this posterior face-selectivity is a result of top-down attentional effects previously reported during face-viewing (Mo et al., 2018). Following fusiform responses, face-selectivity was then seen in more anterior VTC.

While delays in processing along the posterior-to-anterior VTC axis for both faces and words is somewhat consistent with feedforward models of visual processing, the relative latencies are far longer than would be expected in these models (Thorpe et al., 1996; Kravitz et al., 2013). These results instead suggest more extended dynamics, perhaps governed by recurrent processes (Kravitz et al., 2013), with different category-selective regions contributing differentially to multiple, temporally extended stages of face- and word-processing (Ghuman et al., 2014; Hirshorn et al., 2016; Li et al., 2018). Further studies are required to identify these stages and link them to different spatiotemporal patterns of VTC activity. It is important to acknowledge that when analyzing the data at this fine granularity, between-participant variability in neural organization may influence the differences observed in dynamics across regions (Zhen et al., 2015; Gao et al., 2018).

The high-resolution maps of category-selectivity within VTC provided here suggest that in addition to more extensively studied word-selective patches within the occipitotemporal sulcus, additional patches of word-selectivity exist along the mid and anterior fusiform gyrus. These patches of word-selectivity differ in their temporal dynamics from word-selective patches along the occipitotemporal sulcus, but still contain information about word identity. How these word-selective regions differentially contribute to reading and the factors that lead to the development of adjoining patches of word- and face-selective regions remain as important outstanding questions. Understanding this complex and dynamic map of selectivity in VTC is necessary to fully understand the organizational and computational principles governing object recognition.

## Conflict of interest statement

The authors declare no competing financial interests.

## Acknowledgements

We would like to thank the patients, their families, and the clinical staff at the Epilepsy Monitoring Unit at the University of Pittsburgh Medical Center, without whom this study would not be possible. We would also like to thank Marlene Behrmann, David Plaut, Michael Tarr, and the anonymous reviewers for helpful suggestions regarding the analysis and interpretations of the intracranial EEG results; Sean Walls, Ellyanna Kessler, Roma Konecky, and Ashley Whiteman for assistance in intracranial EEG data collection. This study was supported by the National Institutes of Health under R01MH107797 and R21EY030297 and National Science Foundation under 1734907 to A.S.G., National Institutes of Health under T32NS007433-20 to M.J.B., and National Institutes of Health under ZIAMH002909 to C.I.B. and E.H.S.

## Notes

### Competing Interest Statement

The authors have declared no competing interest.

### Summary of Updates

This version of the manuscript has been revised to include additional analyses examining the responses of medial and fusiform word-selective regions, increase the clarity of the methods, and address several additional points of discussion.

